# Phenotypic Variation Reveals Contrasting Ecological Strategies in Wood-Decay Fungal Ecotypes Across a Hybrid Zone

**DOI:** 10.64898/2026.04.09.717513

**Authors:** Ingvild Myhre Ekeberg, Håvard Kauserud, Inger Skrede

## Abstract

Fungi play central roles in terrestrial ecosystem functioning and are the major decomposers of dead wood in forest systems. Wood decay fungi are adapted to growth and decay under different environmental conditions, but we have limited insight into the intraspecific variability in fungal growth and decay. In Fennoscandia, there are two genetically and ecologically distinct ecotypes of the wood-decay fungus *Meruliopsis taxicola*: a northern Continental ecotype associated with Norway spruce (*Picea abies*) growing in moist old-growth forests, and a southern Coastal ecotype growing on Scots pine (*Pinus sylvestris*) in harsher habitats. The two ecotypes hybridize in a narrow contact zone running through Fennoscandia. Here, we investigate the level of adaptation the two ecotypes show in phenotypic traits, and how hybrid isolates perform as compared with the parental genotypes. We performed *in vitro* experiments to quantify mycelial growth rate under varying temperature and drought conditions, as well as decomposition of the two substrates, Scots pine and Norway spruce. Isolates of the Continental ecotype exhibited generally higher growth rates in all environments and caused higher mass loss of both substrates. This is consistent with a more competitive life history strategy in the Continental ecotype, whereas Coastal isolates showed adaptions indicative of greater stress tolerance. Hybrid isolates displayed largely intermediate growth responses relative to the parental ecotypes. Together, these results reveal clear phenotypic divergence between *M. taxicola* ecotypes consistent with contrasting life-history strategies.

## 1 Introduction

Fungi are major components of terrestrial ecosystems in terms of both biomass and diversity and play central roles in ecosystem functioning, particularly through decomposition and nutrient cycling (Crowther et al. 2014; Treseder and Lennon 2015). Understanding how fungi adapt to their habitats is important for understanding their responses to environmental change.

Temperature and moisture are among the most important environmental regulators of wood-decay fungal activity (Lustenhouwer et al. 2020). Wood-decay fungal species show different growth response curves across these variables, mirroring their ecological niches and geographic distributions (Maynard et al. 2019). Most studies documenting variation in growth response curves have focused on interspecific comparisons, whereas intraspecific trait variation and local adaptation in wood-decay fungi remain under-studied. Evidence for intraspecific local adaptation has been reported in fungal plant-pathogens (Laine 2008; Zhan and McDonald 2011), in the model fungus *Neurospora crassa* (Ellison et al. 2011) and in the wild yeast *Saccharomyces paradoxus* (Leducq et al. 2014), suggesting that similar processes may occur in other fungal species. A recent study of the wide-spread wood-decay fungus *Fomitopsis pinicola* in Norway, however, found limited support for phenotypic variability in growth and local adaptation (Kauserud et al. 2024). This indicates that the extent and ecological significance of such variation remain unresolved.

Compared to animals and plants, trait-based approaches have historically been overlooked in fungal ecology (Aguilar-Trigueros et al. 2015; Crowther et al. 2014), probably because it is more difficult to obtain trait data for fungi. However, there are more recent efforts to implement trait-based analyses in the study of fungal life-history strategies (Crowther et al. 2014; Maynard et al. 2019). The classical C-S-R framework, originally describing how plants adapt to three main strategies (competitive (C), stress-tolerant (S) or ruderal (R)) in different environments (Grime 1977), has been adapted to fungi to understand continuous variation in physiological traits linked to species distributions and ecosystem functioning (Crowther et al. 2014; Maynard et al. 2019). In wood-decay fungi, a trade-off between competitive ability and stress tolerance has been recognized as a key axis of life-history variation, where some individuals exhibit rapid mycelial growth that enhances competitive ability under favorable conditions, whereas others grow more slowly but invest more in stress-related traits in harsher environments (Crowther et al. 2014; Lustenhouwer et al. 2020; Maynard et al. 2019). Intraspecific variation in such traits may thus represent local adaptations, where a lineage is locally adapted if it shows higher fitness in its habitat compared to other lineages (Kawecki and Ebert 2004).

Here, we use the wood-decay fungus *Meruliopsis taxicola* to investigate lineage-specific local adaptation in growth and host substrate across a naturally occurring hybrid zone in Fennoscandia. Two genetically and ecologically separated *M. taxicola* ecotypes occur in this region with distinct geographic distributions (Fig. 1): a Coastal ecotype and a Continental ecotype (Ekeberg et al. 2026; Kauserud, Hofton and Sætre 2007). The Coastal ecotype is found in southern areas on dry, exposed logs and branches of Scots pine (*Pinus sylvestris*) in arid, coastal habitats, where it likely grows under relatively stressful conditions due to high temperatures during the growing season and limited water availability. In contrast, the Continental ecotype is distributed in more northern areas on logs of Norway spruce (*Picea abies*) in moist old-growth forests, probably under more favorable growing conditions. The two ecotypes likely diverged in different glacial refugia during the last glaciation, followed by post-glacial migration with their respective hosts into Fennoscandia, before they met in secondary contact and a narrow hybrid zone running through Fennoscandia was established (Ekeberg et al. 2026). Mating between the divergent lineages may lead to different evolutionary outcomes. Hybrid individuals may exhibit trait values similar to one parental lineage, intermediate between the parental lineages, or outside the parental range, depending on the genetic architecture of the underlying traits. The persistence of hybrid individuals is contingent on how trait values interact with the local environment, by determining the fitness of the hybrids relative to the parental lineages.

**Figure 1.**
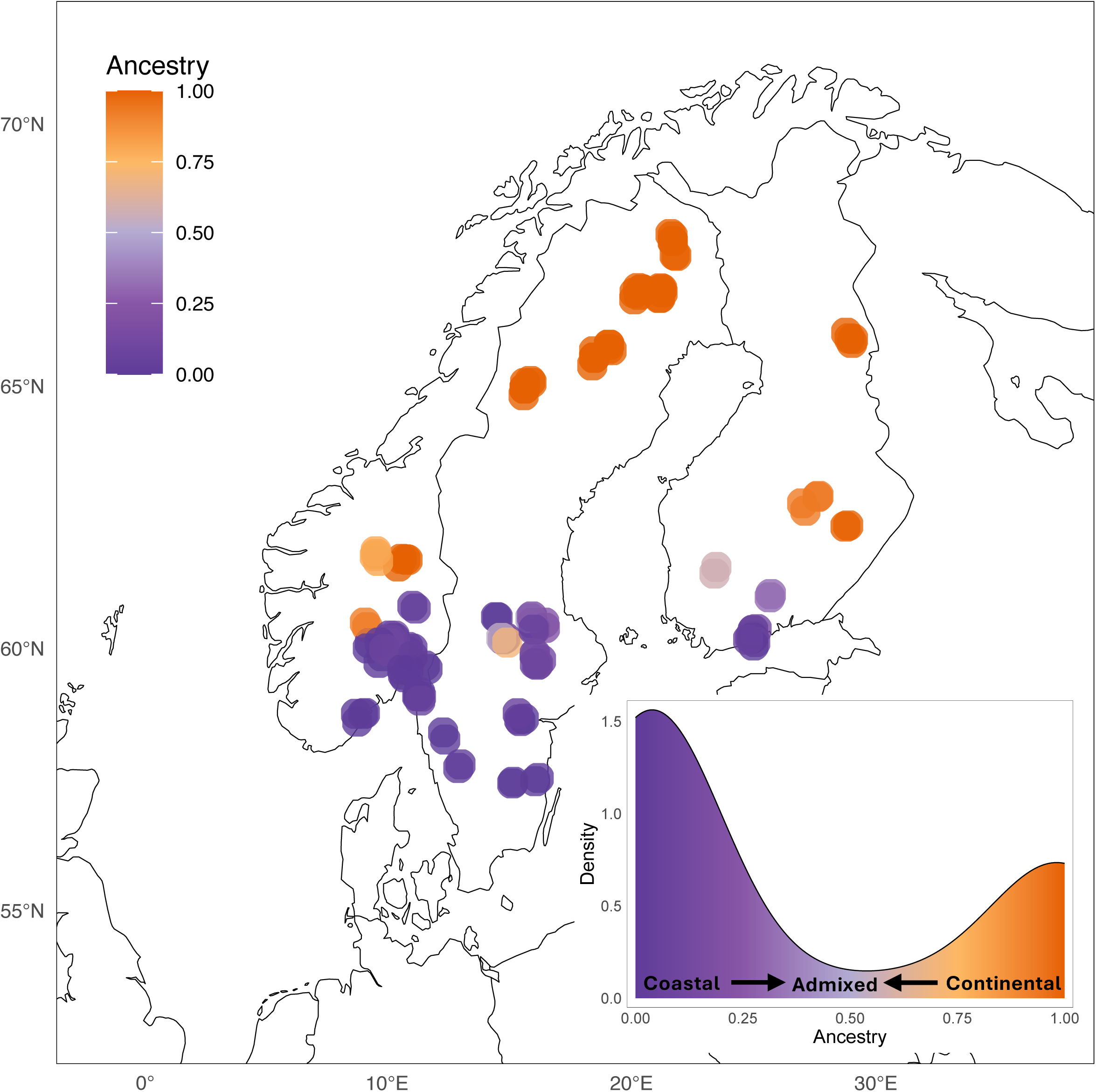
Distribution of *Meruliopsis taxicola* isolates in Fennoscandia. Points are colored by genetic ancestry based on the genetic analyses in Ekeberg et al. (2026), where 0 corresponds to the Coastal ecotype (purple) and 1 to the Continental (orange) ecotype; intermediate values indicate varying degrees of genetic ancestry. The plot on the bottom right shows the density of the ancestry proportion for the isolates included in the study.

The persistence of genetic distinctiveness of the ecotypes despite long-term contact suggests that divergence is sustained by local adaptation. Furthermore, how the hybrid zone is maintained remains unelucidated. Building on the genomic and ecological separation identified in Ekeberg et al. (2026), the aim of this study was to test the level of phenotypic divergence among the ecotypes and their hybrids.

We first ask whether phenotypic growth responses to environmental conditions reflect the ecologies and different distributions of the ecotypes, indicating local adaptations. We hypothesize that the Continental ecotype exhibits faster growth rates compared to the Coastal ecotype across temperature and drought gradients (H1). This prediction is consistent with a more competitive life-history strategy in the Continental ecotype and a more stress-tolerant strategy with slow growth in the Coastal ecotype. Next, we hypothesize that decay will be higher on the ecotype’s preferred substrate (H2), reflecting adaptations to different host tree species. Expected outcomes of this is that the Continental ecotype will have relatively higher decay of Norway spruce while the Coastal ecotype will have higher decay of Scots pine. We further hypothesize that, due to local adaptation, the climate where the isolates were collected explains variation in growth and decomposition responses beyond that attributable to the genetic contribution of either ecotype (H3). We also evaluate phenotypic responses of hybrid isolates with admixed genetic ancestry (i.e. with genetic contribution from both parental lineages) as compared to isolates representing the two parental ecotypes. We may here expect that hybrid growth and decomposition responses are intermediate of the parental ecotypes, reflecting the genetic contribution from the two ecotypes (H4.1). Alternatively, hybrid isolates show low growth and decomposition ability compared to the parental ecotypes (H4.2), due to genetic incompatibilities among the two ecotypes. To test these hypotheses, we conducted growth experiments of *M. taxicola* isolates under temperature and drought gradients, as well as decomposition experiments of Scots pine and Norway spruce wood blocks.

## 2 Material and Methods

### 2.1 Study Material

We included 93 isolates of *M. taxicola* originating from locations spanning a broad climatic gradient across Fennoscandia (Ekeberg et al. 2026). The isolates represented the Coastal and Continental ecotypes, as well as several isolates with hybrid genomic ancestry (i.e. with genetic contributions from both parental ecotypes; Fig 1; Table S1). Genomic ancestry (hereafter “ancestry”) was quantified as admixture proportions inferred under K = 2 genetic clusters and treated as a continuous variable ranging from 0 (≈ Coastal ancestry) to 1 (≈ Continental ancestry), where hybrids are genomic mosaics along this gradient. Details on culturing procedures and the inference of genetic grouping are provided in (Ekeberg et al. 2026). Some of the cultures died during the experiments and were therefore not included in all analyses. The geographic distribution and ancestry of the isolates were visualized on a map of Fennoscandia using the R packages rnaturaleart 1.1.0 (Massicotte and South 2025).

### 2.2 Growth Experiments

We conducted growth (mycelial extension) experiments to compare responses of isolates with differing ancestry. Growth was measured under two different schemes: by manipulating the temperature and the water availability (drought experiment). Each isolate was replicated four times per treatment to account for within-isolate variability and increase statistical power.

Growth rate (mm day^-1^) was calculated for all replicates. Our aim was not to construct complete response curves but to compare growth under optimal conditions representative of those in Fennoscandia. A pilot study indicated negligible growth at 5 °C and 35 °C, consistent with reported optima for other wood-decay Basidiomycetes (Kauserud et al. 2024; Maynard et al. 2019). We therefore selected 20 °C, 25 °C and 30 °C as treatments in the temperature experiment.

Mycelial plugs of standardized size were cut from established cultures using the base of a 1000 µL pipette tip (8 mm in diameter) and placed in the center of 3% malt extract agar (MEA) plates. Plates were incubated at 20 °C in the dark for two days to initiate growth. Two reference marks were drawn on either side of the growing mycelia before plates were transferred to incubators. For the temperature experiment, the incubators were set to the treatment temperatures. Measurements were taken between the reference marks and the hyphal tip with a digital caliper once the hyphae approached the end of the petri dish (∼0.5 cm from the edge, Fig. S1). The two measurements per plate were averaged to account for within-replicate variation in growth. Growth rate was obtained by dividing the distance grown by the number of days it took for the culture to reach the end of the petri dish.

For the drought experiments, water availability was manipulated by adding potassium chloride (KCl) to the medium to achieve three water potential (MPa) treatments. We used the water potentials-0.5 MPa (control; 0 g KCl/L 3% MEA),-1.0 MPa (“dry”; 9 g KCl/L 3% MEA) and-1.5 MPs (“dryest”; 18 g KCl/L 3% MEA) which have been reported to cover growth optima in other wood-decay Basidiomycetes (Kauserud et al. 2024; Maynard et al. 2019). The relationship between water potential and KCl was established in previous studies (Jones, Stewart and Whipps 2011; Maynard et al. 2019; Whiting, Khan and Gubler 2001).

Plates were placed in airtight boxes containing buffer solutions at the bottom of the boxes with matching KCl concentrations to minimize fluctuations in water potential (Maurice et al. 2011; Sautour et al. 2001). Initiation, inoculation and growth measurements followed the same procedure as in the temperature experiment, except that all incubation were conducted in the airtight boxes placed in 20 °C incubators in the dark.

### 2.3 Wood Decomposition Experiments

We tested whether lineages of *M. taxicola* show substrate specialization by quantifying mass loss of inoculated Scots pine and Norway spruce wood blocks in a decomposition experiment. Each isolate was replicated four times per substrate. Wood blocks of the dimension 2 cm x 3 cm x 0.25 cm were obtained from logs of Scots pine and Norway spruce. The blocks were cut such that the shortest dimension (0.25 cm) was perpendicular to the xylem, avoiding heartwood. Blocks were dried for three days at 70 °C and then weighed prior to inoculation. Mycelial plugs were prepared as described for the growth experiments and placed on rewetted wood blocks on plates containing Czapek dox medium without sucrose (1.1 g/L L-glutamic acid monosodium salt monohydrate, 1 g/L potassium phosphate monobasic, 0.5 g/L magnesium sulphate heptahydrate, 20 g/L agar). Nitex nylon mesh (Sefar AG, Heiden, Switzerland) was placed between the wood block and media to prevent mycelium growing into the agar (Fig. S2). Four negative controls (no fungal inoculum) per substrate were included. Plates were incubated in the dark at 20 °C for 125-129 days.

After incubation, mycelium was removed, and wood blocks were dried and weighed again to obtain percentage mass loss. Controls exhibited small mass loss changes (pine: mean 4.72% ± 1.71; spruce: 3.15% ± 0.28), likely reflecting natural variability in wood block mass due to factors such as fluctuations in moisture content and minor handling-related mass loss. Such variation is expected across all blocks and cannot be fully separated from fungal decomposition. Mass loss estimates were therefore interpreted cautiously.

### 2.4 Statistics

We assessed associations between experimental responses and ancestry using linear models. Percentage mass loss was log-transformed to improve normality (Fig. S3). To evaluate relationships among responses and ancestry, and to identify potential trade-offs, we calculated correlations between mean isolate responses (averaged across replicates) in each treatment and visualized the correlation matrix using the corrplot 0.95 (Wei and Simko 2024) package in R. Additionally, we tested differences in mean log-transformed mass loss of pine and spruce wood-blocks with Welch’s two-sample t-test.

Linear mixed effect (LME) models (Pinheiro and Bates 2000) were used to evaluate the effects of treatments, ancestry and climate on growth rate in the temperature and drought experiments and on log-transformed mass loss in the decomposition experiment (negative values were removed). The 19 bioclimatic variables were downloaded from WorldClim 2.1 (Fick and Hijmans 2017) using the geodata 0.6-6 (Hijmans 2025a) R package at 2.5‘ resolution. A PCA of the scaled climatic variables at isolate locations and climatic vectors associated with PC1 and PC2 were identified using the R packages terra 1.8-86 (Hijmans 2025b) and factoextra 1.0.7 (Kassambra and Mundt 2020). Bioclim1 (annual mean temperature) and Bioclim13 (precipitation of wettest month) were selected for LME models because they were weakly correlated with each other (Fig. 2) and had high loadings on PC1 and PC2 (Fig. 2, Table S2). Because climate, geography and genetic structure are confounded, models including the climatic predictors were interpreted cautiously.

**Figure 2.**
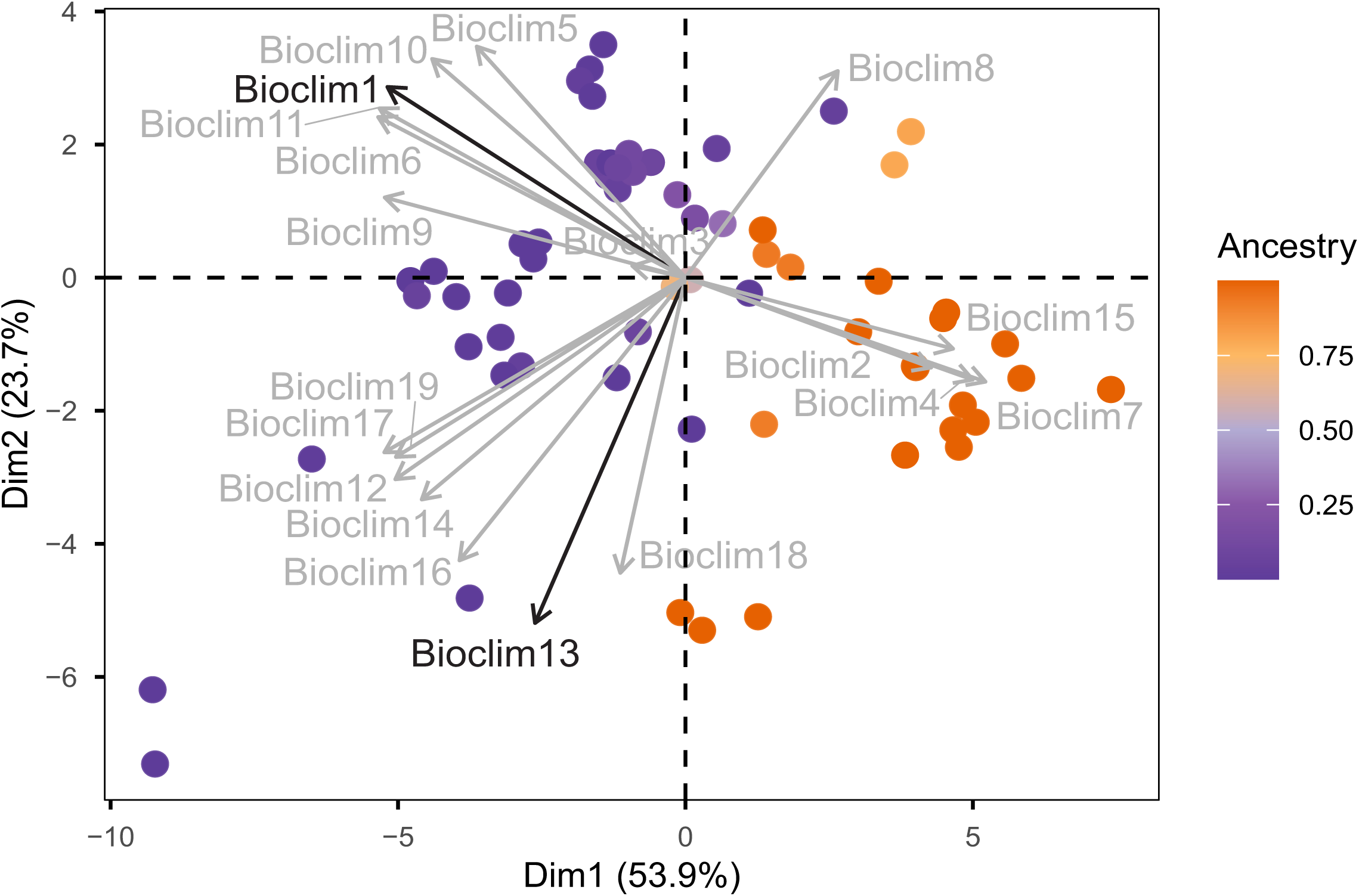
PCA biplot of the 19 Bioclimatic variables in the geographic sampling locations (latitude and longitude) of *Meruliopsis taxicola* in Fennoscandia. Points are colored by genetic ancestry based on the genetic analyses in Ekeberg et al. (2026), where 0 corresponds to the Coastal ecotype (purple) and 1 to the Continental (orange) ecotype; intermediate values indicate varying degrees of genetic ancestry. The Bioclimatic variables are shown as vectors, where Bioclim1 and Bioclim13 have the highest, uncorrelated PC loadings and are highlighted in black.

All models included replicate nested within isolate as a random effect. Heteroscedasticity among treatment levels was addressed by incorporating variance structures (weights) in the models. To determine the fixed effect structure, we fitted candidate models using maximum likelihood in the nlme 3.1-168 (Pinheiro and Bates 2000; Pinheiro, Bates and R Core Team 2025) R package. Model selection was based on Akaike’s Information Criterion (AIC).

Following Burnham and Anderson (2004) rules of thumb, models with Δ AIC = 0 had the highest support, while models Δ AIC ≤ 2 were considered to have substantial/equal support, those with 4 ≤ Δ AIC ≤ 7 to have considerably less support, and those with Δ AIC > 10 to have no support. Candidate models were also compared using likelihood ratio tests (ANOVA; Table S3). When comparing several equally supported models, we chose the most parsimonious one, which was refitted using restricted maximum likelihood (REML). The refitted models were evaluated with ANOVA to evaluate the contributions of each fixed effect to the variance in the response. Model assumptions were evaluated with residual diagnostics (Fig. S4) using the R package predictmeans 1.1.1 (Luo 2024). Random effects were inspected using the ranef function in the nlme pacakge to extract the random effects (Fig. S5).

For each selected model refitted using REML, we evaluated the effect of ancestry on the response variable within each treatment. Slopes were estimated using the emtrends function in the emmeans 2.0.1 (Lenth and Piaskowski 2025) R package. Pairwise differences among the slopes were assessed using model-based Wald t-tests, where p-values were Tukey corrected when there were more than two comparisons (i.e. the temperature and drought experiments). To visualize the interaction between treatment and ancestry, we obtained model-predicted marginal means across 100 evenly spaced ancestry values (0-1) for each treatment using the emmeans function. The predicted responses were plotted with 95% confidence levels.

### 2.5 Data Manipulation and Visualization

Data was manipulated and visualized with the tidyverse 2.0.0 package (Wickham 2016) in R, if not stated otherwise. Final graphic adjustments of figures were made in Microsoft PowerPoint 16.102.2 (Microsoft Corporation, 2025).

## 3 Results

### 3.1 Growth Rate and Mass Loss Vary With Genomic Ancestry and Environmental Conditions

Growth rates of the isolates increased from isolates with a fully Coastal ecotype ancestry to Continental ecotype ancestry across all temperature treatments. Thus, the Continental genotypes exhibited faster growth across all temperature treatments (Fig. 3A, Table 1).

**Figure 3.**
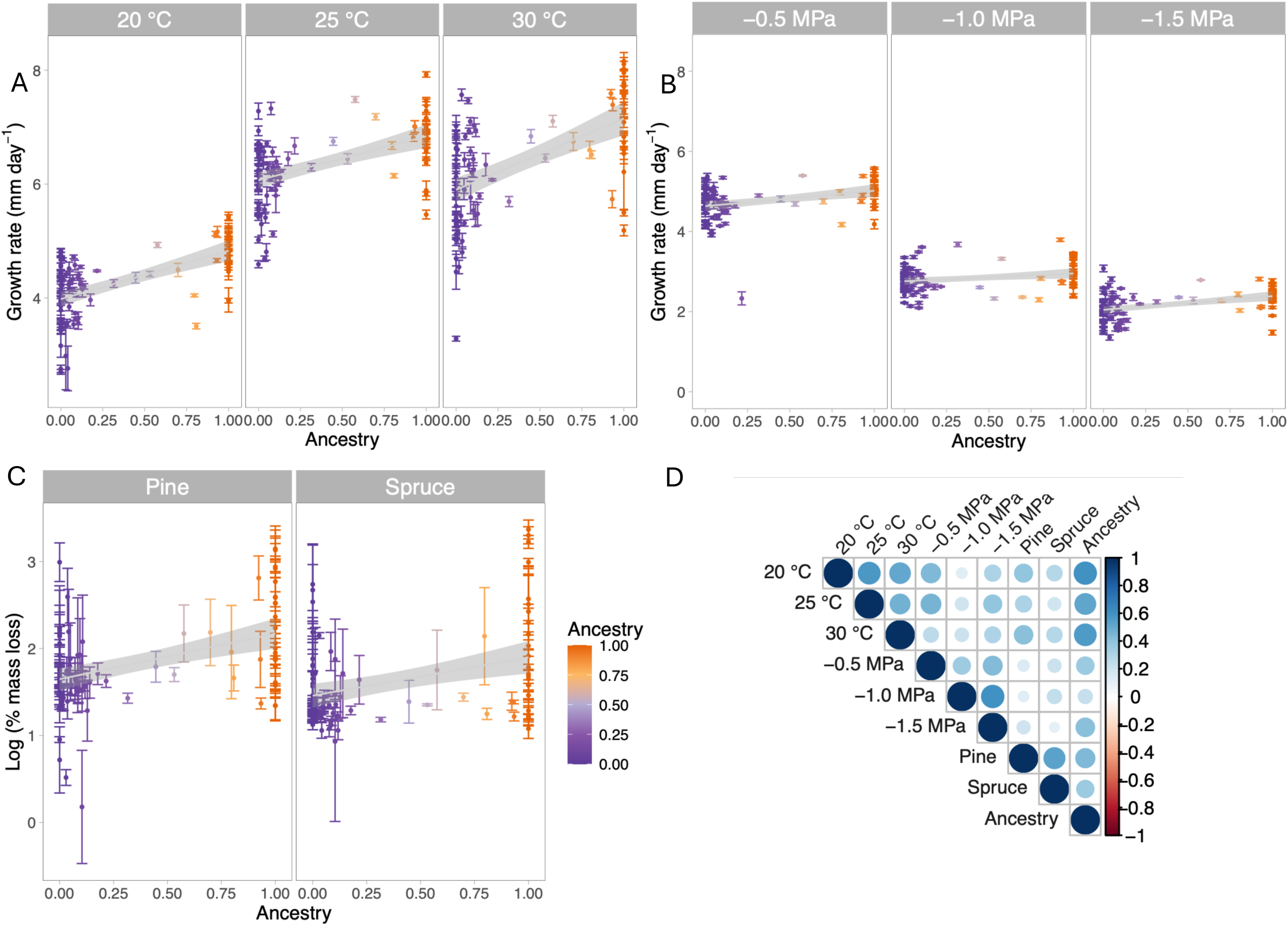
Relationship between phenotypic responses of *Meruliopsis taxicola* isolates and genetic ancestry. A-C) Mean response (± SE) across replicates within each treatment plotted against ancestry. Grey shaded area represents 95% confidence intervals from fitted linear regression models. Points are colored by ancestry, where 0 corresponds to the Coastal ecotype (purple) and 1 to the Continental ecotype (orange); intermediate values indicate varying degrees of ancestry. A) Growth rate (mm day^-1^) under three temperature treatments (20 °C, 25 °C and 30 °C; n = 91). B) Growth rate (mm day^-1^) under three drought (water potential) treatments (-0.5 MPa,-1.0 MPa and-1.5 MPa; n = 89). C) Decomposition responses measured as log-transformed percentage gram mass loss of pine or spruce wood blocks. D) Correlation matrix (Pearson’s correlation coefficient) of mean responses across all treatments shown in panels A-C. Circle size and color denote the strength and direction of correlations.

**Table 1.**
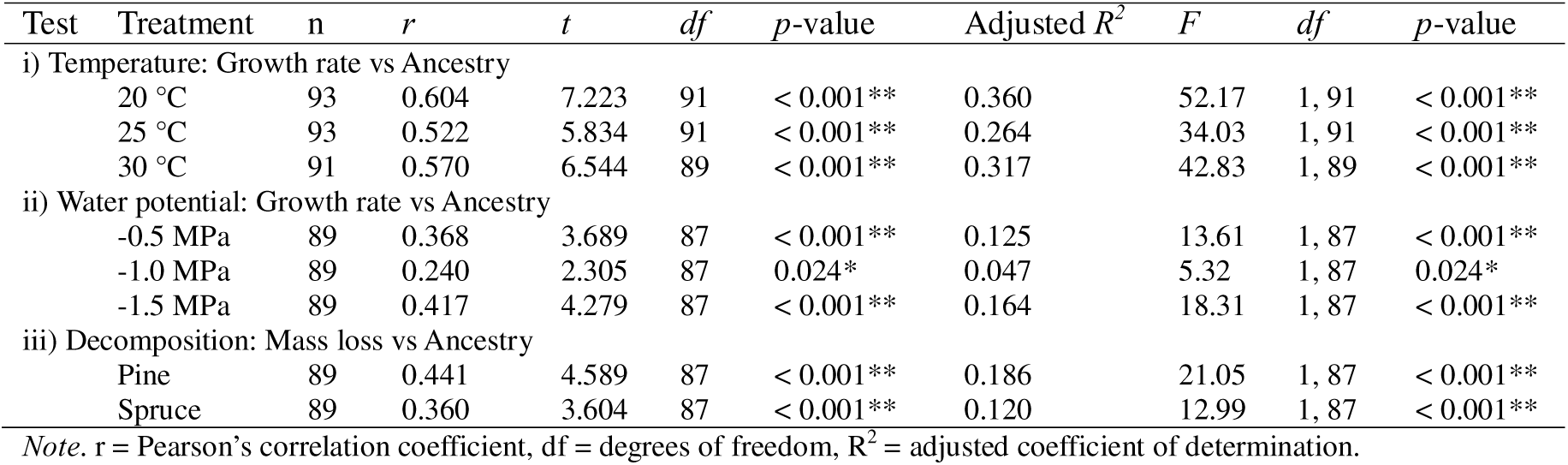
Relationship between responses in the temperature, drought and decomposition experiments and ancestry proportion. Correlation tests and linear regression models for each treatment are shown. Responses are measured as the mean across replicates for each isolate. The response in the temperature and water potential experiments is measured in growth rate per day (mm day^-1^) and the response in the decomposition experiment is given as log mass loss of wood blocks (%).

However, there was considerable variation among isolates of both ecotypes, suggesting that the ecotype category only accounted for parts of the phenotypic variation. Notably, growth rate was highly consistent across replicates, demonstrating that the growth responses reflect inherent biological differences rather than methodological bias. Several linear mixed effect (LME) models were considered equally fit in explaining growth rate under different temperatures (i.e. models i.3, i.6 and i.7 with Δ AIC < 2; Table 2). The most parsimonious of these models included only the interaction term between temperature treatment and genomic ancestry (model i.3) and was selected. ANOVA of the refitted model showed that temperature treatment and genomic ancestry had an overall strong effect on growth rate, with a significant interaction term (*p* < 0.01; Table 3). Hence, the effect of level of genomic ancestry from the two ecotypes varied within each temperature treatment. In particular, the predicted positive effect on growth rate of high levels of continental ecotype ancestry was stronger at higher temperatures (Fig. 4A, Table 4-5), suggesting that the Continental ecotype grows relatively faster in warmer temperatures compared to the Coastal ecotype.

**Figure 4.**
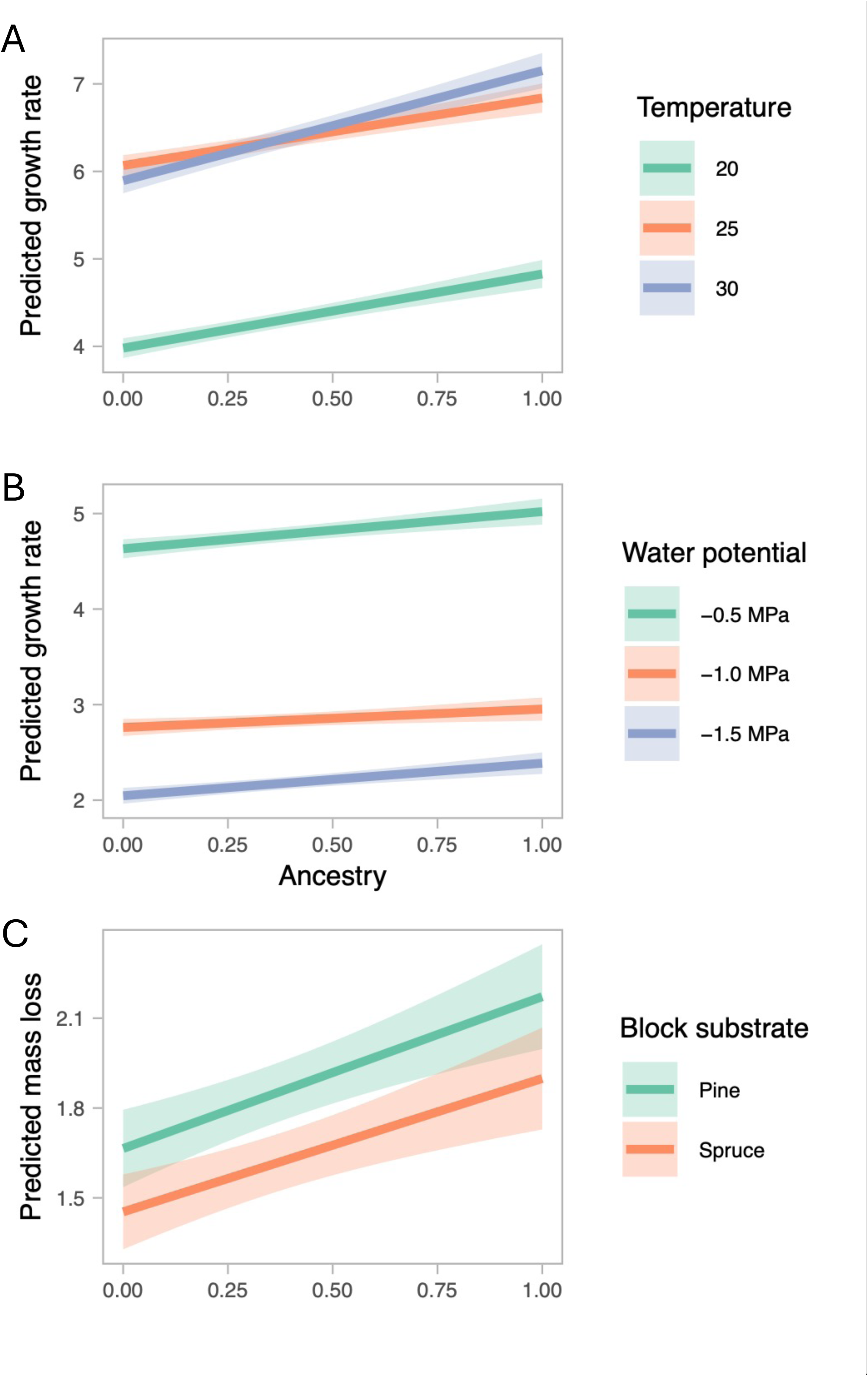
Predicted marginal mean responses for *Meruliopsis taxicola* in A) temperature, B) drought (water potential) and C) decomposition experiments. Responses are plotted against admixture proportions where 0 corresponds to the Coastal ecotype and 1 to the Continental ecotype; intermediate values indicate varying degrees of ancestry. Marginal means were calculated using emmeans in R from the refitted REML linear mixed effect models, including an interaction term between ancestry and treatment, for 100 evenly spread values of the ancestry variable. Solid lines show the predicted responses and shaded area represents 95% confidence intervals for each treatment.

**Table 2.** Comparison of candidate models in linear mixed effect models within each of three experiments (temperature, drought and decomposition) for *Meruliopsis taxicola*, fitted with ML. The hypotheses tested for each candidate model are given (see main text). ΔAIC values show the difference in AIC compared to the candidate model with the lowest AIC within each analysis. Bold text indicates both the best fit and the most parsimonious model refitted in downstream analyses. Equivalently best-fit models with ΔAIC < 0.2 are highlighted in italics. See material and methods and supplementary information for additional details.

| Experiment | Hypothesis | Candidate model | Fixed effects structure | df | AIC | ΔAIC |
| --- | --- | --- | --- | --- | --- | --- |
| **i) Temperature** |  |  |  |  |  |  |
|  | H1 | i.1 | Temp | 10 | 1754.742 | 68.291 |
|  | H1, H4 | i.2 | Temp + Ancestry | 11 | 1705.113 | 18.662 |
|  | H1, H4 | **i.3** | **Temp\*Ancestry** | **14** | **1686.809** | **0.358** |
|  | H1, H3 | i.4 | Temp + Bioclim1 + Bioclim13 | 10 | 1728.459 | 42.008 |
|  | H1, H3 | i.5 | Temp*Bioclim1 + Bioclim13 | 12 | 1711.598 | 25.147 |
|  | H1, H3, H4 | i.6 | Temp*Ancestry + Bioclim1 + Bioclim13 | 13 | 1686.451 | 0.000 |
|  | H1, H3, H4 | i.7 | Temp*Ancestry + Bioclim1*Ancestry + Bioclim13 | 14 | 1687.101 | 0.650 |
| **ii) Drought** |  |  |  |  |  |  |
|  | H1 | ii.1 | WP | 8 | 316.009 | 29.988 |
|  | H1, H4 | ii.2 | WP + Ancestry | 9 | 300.148 | 14.126 |
|  | H1, H4 | **ii.3** | **WP\*Ancestry** | **11** | **286.021** | **0.000** |
|  | H1, H3 | ii.4 | WP + Bioclim1 + Bioclim13 | 10 | 300.632 | 14.611 |
|  | H1, H3 | ii.5 | WP*Bioclim1 + Bioclim13 | 12 | 288.431 | 2.409 |
|  | H1, H3, H4 | ii.6 | WP*Ancestry + Bioclim1 + Bioclim13 | 13 | 288.287 | 2.266 |
|  | H1, H3, H4 | ii.7 | WP*Ancestry + Bioclim1*Ancestry + Bioclim13 | 14 | 288.437 | 3.416 |
| **iii) Decomposition** |  |  |  |  |  |  |
|  | H2 | iii.1 | Substrate | 6 | 1378.883 | 18.039 |
|  | H2, H4 | iii.2 | Substrate + Ancestry | 7 | 1361.052 | 0.209 |
|  | H2, H4 | **iii.3** | **Substrate\*Ancestry** | **8** | **1362.675** | **1.832** |
|  | H2, H3 | iii.4 | Substrate + Bioclim1 + Bioclim13 | 8 | 1360.843 | 0.000 |
|  | H2, H3 | iii.5 | Substrate*Bioclim1 + Bioclim13 | 9 | 1362.295 | 1.452 |
|  | H2, H3, H4 | iii.6 | Substrate*Ancestry + Bioclim1 + Bioclim13 | 10 | 1364.334 | 3.491 |
|  | H2, H3, H4 | iii.7 | Substrate*Ancestry + Bioclim1*Ancestry + Bioclim13 | 11 | 1361.840 | 0.997 |

Note. Temp = Temperature, WP = water potential, df = degrees of freedom, AIC = Akaike Information Criterion, \* = interaction term.

**Table 3.** ANOVA table for the selected model (see Table 1 for model details) for each experiment (i. temperature, ii. drought and iii. decomposition), refitted with REML.

| Model | Effect | numDF | denDF | *F*-value | *p*-value |
| --- | --- | --- | --- | --- | --- |
| i.3 | Intercept | 1 | 731 | 4868.075 | < 0.0001** |
|  | Temp | 2 | 731 | 1958.412 | < 0.0001** |
|  | Ancestry | 1 | 91 | 67.847 | < 0.0001** |
|  | Temp:Ancestry | 2 | 731 | 11.432 | < 0.0001** |
| ii.3 | Intercept | 1 | 707 | 8467.944 | < 0.0001** |
|  | WP | 2 | 707 | 4002.362 | < 0.0001** |
|  | Ancestry | 1 | 87 | 19.153 | < 0.0001** |
|  | WP:Ancestry | 2 | 707 | 9.228 | < 0.0001** |
| iii.2 | Intercept | 1 | 611 | 655.972 | < 0.0001** |
|  | Substrate | 1 | 611 | 13.593 | < 0.0002** |
|  | Ancestry | 1 | 87 | 19.640 | < 0.0001** |
|  | Substrate:Ancestry | 1 | 611 | 0.375 | 0.5405 |

*Note.* numDF = Numerator degrees of freedom, denDF = Denominator degrees of freedom, WP = Water Potential, Temp = Temperature.

**Table 4.** Pairwise tests of differences in estimated slopes withi calculated from the REML models using the emtrends function in R. All REML models included an interaction term between treatment and ancestry. The estimate is the difference in slopes for each comparison of treatments (contrast). *p*-values were adjusted with the Tukey method when more than three treatments were compared (i.e. the temperature and drought experiments).

| Experiment | Contrast | Estimate | SE | *df* | *t* | *p*-value |
| --- | --- | --- | --- | --- | --- | --- |
| **Temperature** | 20 °C – 25 °C | 0.077 | 0.064 | 731 | 1.198 | 0.455 |
|  | 20 °C – 30 °C | -0.409 | 0.100 | 731 | -4.191 | < 0.0001 |
|  | 25 °C – 30 °C | -0.486 | 0.103 | 731 | -4.731 | < 0.0001 |
| **Drought** | -0.5 MPa – -1.0 MPa | 0.200 | 0.061 | 707 | 3.198 | 0.0041 |
|  | -0.5 MPa – -1.5 MPa | 0.048 | 0.053 | 707 | 0.893 | 0.6447 |
|  | -1.0 MPa – -1.5 MPa | -0.149 | 0.036 | 707 | -4.089 | 0.0001 |
| **Decomposition** |  |  |  |  |  |  |
|  | Pine – Spruce | 0.062 | 0.101 | 611 | 0.612 | 0.5405 |

Note. *df* = degrees of freedom, MPa = water potential.

**Table 5.** Predicted response slopes from REML linear mixed effect models within treatment for each experiment, estimated with emmeans in R. The models included an interaction term between treatment and ancestry. 95% confidence levels for the estimates are shown.

| Experiment | Treatment | Ancestry trend | SE | df | Lower CL | Upper CL |
| --- | --- | --- | --- | --- | --- | --- |
| **Temperature** |  |  |  |  |  |  |
|  | 20 °C | 0.848 | 0.103 | 91 | 0.644 | 1.053 |
|  | 25 °C | 0.771 | 0.108 | 91 | 0.557 | 0.986 |
|  | 30 °C | 1.258 | 0.131 | 91 | 0.998 | 1.517 |
| **Drought** |  |  |  |  |  |  |
|  | -0.5 MPa | 0.389 | 0.089 | 87 | 0.212 | 0.565 |
|  | -1.0 MPa | 0.193 | 0.080 | 87 | 0.034 | 0.351 |
|  | -1.5 MPa | 0.341 | 0.074 | 87 | 0.195 | 0.488 |
| **Decomposition** |  |  |  |  |  |  |
|  | Pine | 0.507 | 0.114 | 87 | 0.280 | 0.734 |
|  | Spruce | 0.445 | 0.111 | 87 | 0.225 | 0.666 |

Note. MPa = water potential of the medium, SE = standard error, df = degrees of freedom and CL = 95% confidence levels.

There was also a positive relationship between growth rate and genomic ancestry across all three water potential treatments, simulating different drought conditions (Fig. 3B, Table 1). As in the temperature experiment, there was high variability in growth rate among isolates of both ecotypes, but little variation across replicates, suggesting little methodological bias. The LME model which best explained variation in growth rate under different water potentials, included an interaction term between water potential and genomic ancestry (Table 2).

ANOVA of the refitted model showed that water potential and genomic ancestry explained most of the variation in growth rate (Table 3), as well as a significant interaction effect between them (*p* < 0.01; Table 3). Hence, the effect of drought on growth depends on the genomic contribution from either ecotype (Table 3). The effect of genomic ancestry on predicted growth rate was most pronounced in the control (-0.5 MPa) and the simulated driest (-1.5 MPa) condition (Fig. 4B, Table 4-5), implying that the difference in growth rate between the Continental and Coastal ecotypes is less pronounced under more intermediate drought conditions.

Isolates of Continental ancestry generally caused more mass loss than Coastal isolates in both substrates (Fig. 3C, Table 1). However, as for the previous growth experiments, there was considerable variation between isolates of both ecotypes. Additionally, variation across replicates was considerably higher than in the previous growth experiment, suggesting more methodological bias. Several LME models including genomic ancestry and climatic variables could explain variation in mass loss (Table 2). The simplest model included an interaction between substrate type and genomic ancestry and was chosen for downstream analyses.

However, ANOVA of the refitted model revealed that the interaction was not significant (*p* > 0.01; Table 3). Moreover, predicted effects of ancestry on mass loss were similar between pine and spruce substrates (Fig. 4C), and pairwise comparisons of the estimated slopes did not differ (Table 4). These results support that the effect of genomic ancestry on mass loss is consistent across substrate types, where the Continental isolates consistently cause higher mass loss compared to the Coastal isolates.

In sum, across all three experiments, the growth and decay rates generally increased along the genomic gradient going from fully Coastal to Continental ancestry (Fig. 3A-C, Table 1), indicating consistently higher performance of the Continental ecotype across environmental gradients. Only positive relationships were detected when comparing the average growth responses of isolates across the three experiments (Fig. 3D).

### 3.2 Hybrid Trait Responses Are Intermediate of Parental Ecotypes

Hybrid isolates with admixed genomic backgrounds (∼0 > ancestry < ∼1) of the Coastal and Continental ecotypes typically exhibited intermediate growth and mass loss responses relative to isolates with predominantly Coastal or Continental ancestry. This pattern was largely continuous, with growth rate and mass loss increasing linearly with increasing Continental ancestry (Fig. 3A-C, Table 1). Despite this overall linear trend, substantial variation among isolates was observed in some treatments. Notably, some isolates with intermediate genomic ancestry showed growth rates equal to or exceeding those of Continental isolates, particularly in the two driest drought treatments. Taken together, these results indicate predominantly linear hybrid trait responses, with mostly intermediate growth and mass loss responses compared to isolates of fully Coastal or Continental ancestry.

## 4 Discussion

### 4.1 Phenotypic Divergence Suggests Local Adaptation in Two *Meruliopsis taxicola* Ecotypes

Isolates of the Continental ecotype exhibited generally higher growth rates across all treatments in the temperature and drought experiments compared to Coastal isolates, supporting our first hypothesis (H1). However, there was considerable variation between isolates within both ecotypes, indicating high individual variability. Moreover, isolates of the Continental ecotype caused greater mass loss of both pine and spruce wood blocks, though there was considerable between and within-isolate variation. These results did not support our second hypothesis (H2), where we expected host-specific decay responses. Taken together, these results suggest that the Continental ecotype may invest more in rapid radial growth and decay rate, consistent with a competitive life history strategy, whereas the Coastal ecotype may allocate relatively more resources to stress-related traits, resulting in slower growth and decay rates (Lustenhouwer et al. 2020; Maynard et al. 2019). Although we did not measure combative traits in our study, competitive ability has previously been shown to be positively correlated with growth rate in other wood-decay basidiomycetes (Lustenhouwer et al. 2020; Maynard et al. 2019), but it remains to be tested if this holds up the two ecotypes of *M. taxicola*.

Although our observations can be linked to ecotypic differentiation to C-selected and S-selected life-history strategies, we do not have direct assessments of this, and we recognize that there might be other explanations for the differences in growth rate and mass loss. Genomic data suggests that the Coastal ecotype is highly inbred and has relatively low genetic diversity compared to the Continental ecotype (Ekeberg et al. 2026). Inbreeding and inbreeding depression can have substantial effects on phenotypic traits and fitness (Charlesworth and Willis 2009). For example, inbred strains of the Basidiomycete bird’s nest fungus *Cyathus stercoreus* showed significant reduction in growth rates compared to outcrossed specimens (Malloure and James 2013). Similarly, growth rate, among other traits, was reduced in inbred versus outbred laboratory populations of the commercial button mushroom *Agaricus bisporus* (Xu 1995). Thus, it is possible, at least partly, that differences in responses may be due to the inbred nature of the Coastal ecotype, or because of some other unexplored reason.

In the growth experiments (temperature and drought), the selected linear mixed effects (LME) models both included significant interaction terms between genomic ancestry and environmental gradients. This interaction indicates that the isolates growth responses to environmental variation depend on genomic background, which is consistent with reaction norm divergence (Murren et al. 2014), and may reflect local adaptation. The two *M. taxicola* lineages are genomically distinct and distribution modelling has also indicated that the distribution of ecotypes associates with different climatic variables (Ekeberg et al. 2026).

Thus, it is not unexpected *per se* that the ecotypes have diverged also in phenotypic traits and putatively adapted to different environments. Notably, despite ongoing gene flow between the ecotypes and the existence of a hybrid zone, divergence has not been erased during the time they have lived in sympatry, suggesting that differentiation is maintained by local adaptation. However, following the definition by Kawecki and Ebert (2004), fitness of one ecotype in its home environment compared to the environment of the other ecotype should be assessed to conclude whether they are locally adapted. This could be tested in field reciprocal transplant experiments where survival of the ecotypes is quantified, which, for example, has been done in *Saccharomyces paradoxus* (Boynton et al. 2017).

We included climatic variables in our statistical models to evaluate whether climate could explain variation in trait responses beyond Coastal or Continental genomic ancestry (H3). Some models that included climate variables were considered equally good in explaining variance in trait responses as those only including treatment and genomic ancestry. Genomic studies of other fungal species have identified loci associated with climatic variables, e.g. in *Phellopilus nigrolimitatus* (Sønstebø et al. 2022), *N. crassa* (Ellison et al. 2011) and *Suillus brevipes* (Branco et al. 2017), suggesting the potential for climate-mediated adaptive differentiation in fungi. However, in our study, climate, geography and genetic structure are confounded, preventing clear partitioning of their effects. Climatic variables were therefore excluded from the selected models to avoid overparameterization. As a result, our third hypothesis (H3) remains inconclusive.

Despite that Continental isolates on average causes higher mass loss of both substrates than Coastal isolates, the two ecotypes are largely host-specific in nature. Even though the substrates, Norway spruce and Scots pine, grow in sympatry where *M. taxicola* is found, the Continental ecotype prefers Norway spruce while the Coastal type prefers Scots pine. Given the putative competitive life-history strategy of the Continental ecotype, it is therefore interesting that the Continental ecotype in general does not colonizes Scots pine and has not outcompeted the Coastal ecotype and taken over its niche. In this study, we measure decomposition in simplified *in vitro* conditions. However, in nature, dead wood represents heterogenous and dynamic habitats structured by, for example, bark presence, moisture, microbial interactions, and forest structure (Kolényová et al. 2024; Seibold et al. 2021; Ulyshen 2016). Our simplified *in vitro* decomposition experiments obviously do not fully capture the complexity of host conditions in nature. For example, the Continental ecotype is typically found on the bark of moist spruce logs in old-growth forests, while the Coastal ecotype grows on dry, exposed pine wood, often with little bark and on dead branches of living trees (Kauserud, Hofton and Sætre 2007). These contrasting micro-habitats likely impose selective pressures not represented in our laboratory experiment, and may explain the observed host associations. Hence, some unmeasured substrate factor(s) may interact with climate in regulating the host specialization

### 4.2 Intermediate Trait Responses May Explain Hybrid Persistence

Our results largely support the hypothesis that hybrid trait responses depend on the relative genetic contribution from the Coastal and Continental ecotype (H4.1). Across most experimental treatments, trait values increased approximately linearly along the genomic gradient from Coastal to Continental ancestry, where hybrid growth and decomposition responses generally were intermediate relative to the parental ecotypes. Furthermore, we found no support for hypothesis H4.2, which predicted low growth and decomposition responses in hybrid isolates compared to isolates of the parental ecotypes due to genetic incompatibilities.

Even when hybrids overcome genetic incompatibilities (Coyne 1992), and are fertile and viable, intermediate phenotypic traits in hybrids can still lead to reduced fitness in parental habitats (Baack et al. 2015). For instance, the intermediate morphology of the hybrid plant species *Geum intermedium* (*G. rivale* x *Geum urbanum*) is selected against in the natural habitats of its parental species (Ruhsam, Hollingsworth and Ennos 2011; Ruhsam, Hollingsworth and Ennos 2013). In contrast, intermediate phenotypes may also facilitate persistence in environments not occupied by the parental ecotypes or enable exploitation of novel ecological niches (Buggs 2007). For example, hybrids in the seabird genus *Pachyptila,* possess intermediate beak sizes compared to the parental species, which has enabled adaptation to a new feeding niche and homoploid speciation (Masello et al. 2019).

Furthermore, hybrid *M. taxicola* individuals exhibit higher genetic diversity than Coastal specimens (Ekeberg et al. 2026). Such elevated genetic diversity in hybrids may enhance adaptive potential, particularly under changing or heterogenous environmental conditions, due to increased allelic variation available for selection (Vallejo-Marín and Hiscock 2016).

Taken together, the intermediate trait values observed in *M.* t*axicola* hybrids, suggest that they do not have an inherent fitness advantage. The strong ecological adaptations of the Coastal and Continental ecotypes may maintain their separation in ecotype-specific environments. Since hybrids do not show poor performance due to genetic incompatibilities, the *M. taxicola* hybrid zone in Fennoscandia may simply persist within the overlapping distributions of the parental ecotypes. Resolving what sustains the hybrid zone requires direct estimates of hybrid fitness. Future work should therefore quantify hybrid survival and reproductive success, for example by investigating spore establishment and viability and by conducting crossing experiments with hybrid isolates. Additionally, reciprocal transplant experiments of hybrid specimens in environments specific to the parental ecotypes would provide direct assessments of hybrid survival and help elucidate factors sustaining the hybrid zone.

### 4.3 Conclusions and Future Perspectives

Phenotypic trait experiments measuring growth rate under different temperature and drought regimes, as well as decay of pine and spruce wood blocks, revealed divergence in phenotypic traits between the two *M. taxicola* ecotypes. The Continental ecotype exhibited responses consistent with a competitive life-history strategy with fast growth and decay rates. In contrast, the Coastal ecotype displayed responses indicative of a stress-tolerant life-history strategy, with comparatively slower growth and decay rate. These differences possibly reflect adaptations to contrasting climatic conditions, as well as macro and micro-habitats, in Fennoscandia. Responses in hybrid isolates with variable admixed genomic background were generally as expected based on the genomic contribution from the parental ecotypes.

To gain more understanding beyond simplified *in vitro* experiments, future work could include *in situ* experiments assessing growth rate, decay and survival under field conditions. For example, reciprocal transplant decomposition experiments could help clarify the level of natural adaptation and what environmental variables underly growth and decay responses of the ecotypes. In addition, future studies should combine genomic and environmental data to identify candidate genes associated with environmental differentiation. Combining these approaches will provide deeper insight into what maintains the hybrid zone and the divergence between ecotypes.

## Acknowledgements

This work was supported by the Royal Ministry of Education and Research (Norway) and the University of Oslo. We thank Sundy Maurice for the development of laboratory protocols for the water potential experiment. We further acknowledge M. C. Di Luca, R. Falleth, L. G. Gutierrez, C. Mathiesen and G. K. Svendsen for technical assistance, and K. T. Helleland for setting up the decomposition experiment. We thank D. P. Navarro for feedback on preliminary results, and A. K. Brysting for comments on results and on the manuscript.

## Author contributions

IME, HK. and IS conceptualized the study. HK and IS supervised the study. IME performed the experiments, formal analyses, visualizations and data curation. IME wrote the initial draft of the manuscript; HK and IS provided reviews and edits. All authors have read and approved the final version of the manuscript.

## Data availability

Experimental data and records used to produce this study are available at https://dataverse.no/previewurl.xhtml?token=80117771-2f4a-4ab1-8634-b158679e6e98, DataverseNO, DRAFT version.

