## Supplementary figures for "Phenotypic Variation Reveals Contrasting Ecological Strategies in Wood-Decay Fungal Ecotypes Across a Hybrid Zone"

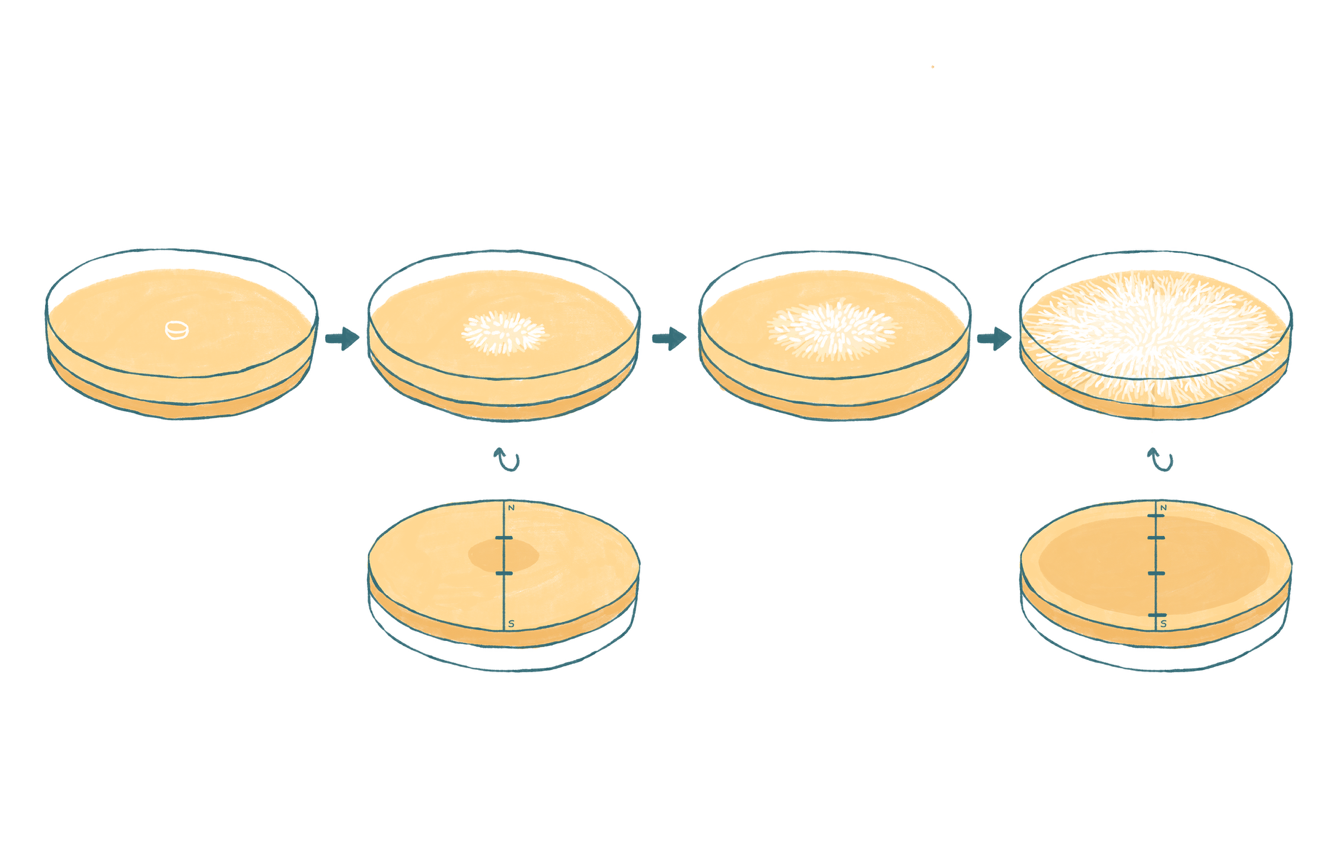

Figure S1
Illustration of the experimental setup for the growth rate experiments. The top row shows petri dishes inoculated with mycelium and radial mycelial extension. The bottom row shows how the underside of the petri dishes and how reference marks are made on the growing mycelium. Growth is measured between the reference marks. Illustration by Marie David.

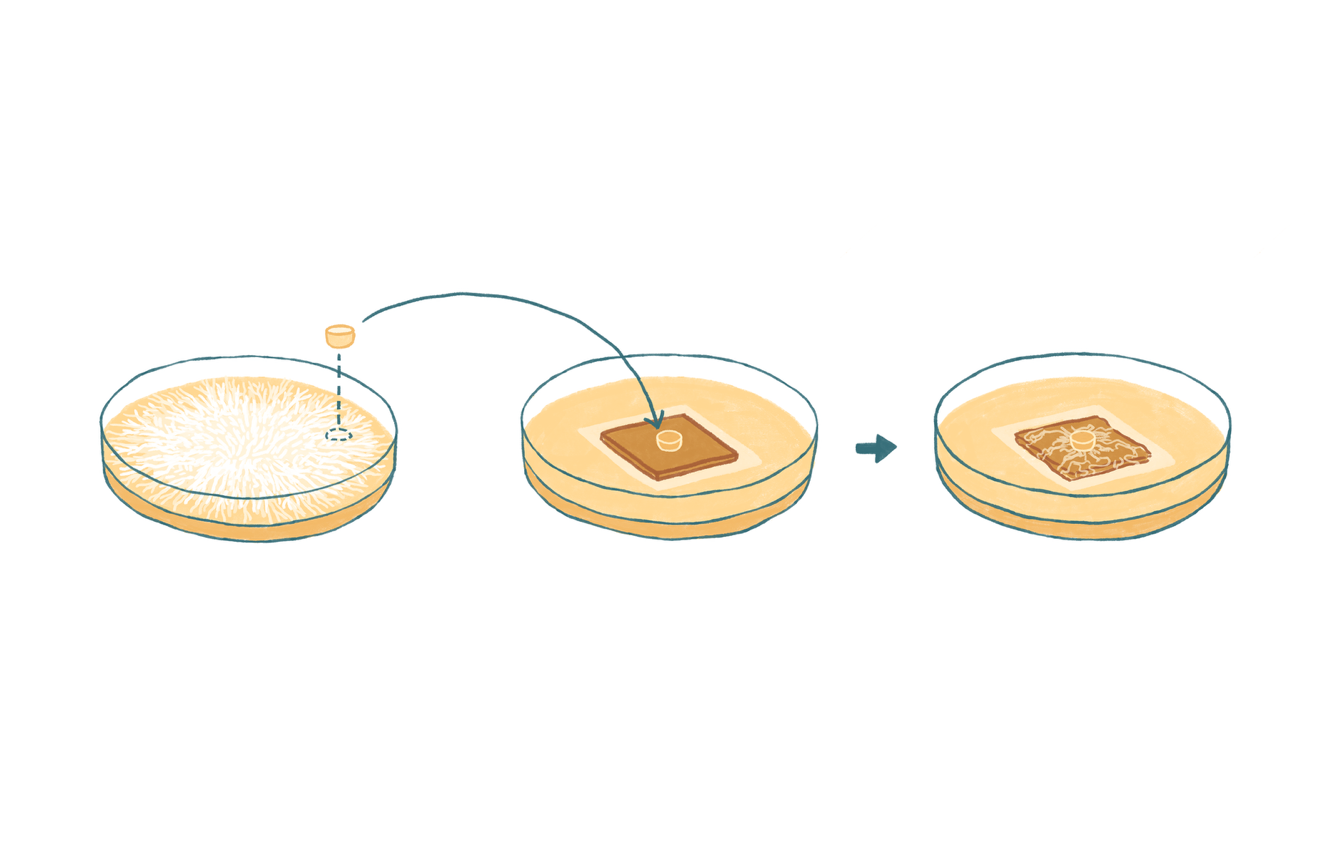

Figure S2
Illustration of the experimental setup for the decomposition experiment. The illustration shows how wood blocks are inoculated by mycelium from established cultures. Wood blocks are separated from the medium with nylon mesh. Illustration by Marie David.

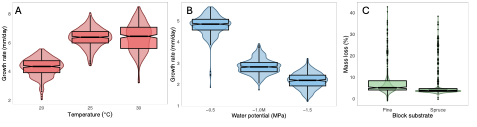

Figure S3
Responses in temperature (A; n = 91), water potential (B; n = 89) and decomposition (C; n = 89) experiments for Meruliopsis taxicola isolates. Combined boxplots and violin plots are shown for each treatment.

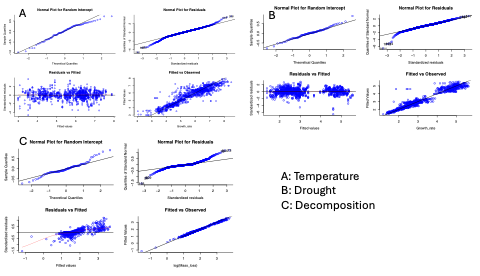

Figure S4
Diagnostic plots for the REML fitted models for the A) temperature, B) drought and C) decomposition experiments. Each panel shows normal plot for random intercept, normal plot for residuals, residuals vs. fitted and fitted vs. observed.

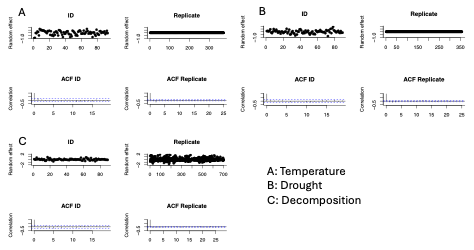

Figure S5
Top panels show random effects in the REML models reflecting variation in samples (cultures) and replicate of the culture for the A) Temperature, B) Drought and C) Decomposition experiment. The bottom panels show autocorrelation functions for the random effects for the A) Temperature, B) Drought and C) Decomposition experiment.
